# A novel tool for designing targeted gene amplicons and an optimised set of primers for high-throughput sequencing in tuberculosis genomic studies

**DOI:** 10.1101/2025.01.13.632698

**Authors:** Linfeng Wang, Naphatcha Thawong, Joseph Thorpe, Matthew Higgins, Mark Tan Kia Ik, Waritta Sawaengdee, Surakameth Mahasirimongkol, Joao Perdigao, Susana Campino, Taane G. Clark, Jody E. Phelan

**Affiliations:** Faculty of Infectious and Tropical Diseases, London School of Hygiene & Tropical Medicine, WC1E 7HT London, UK; Medical Life Sciences Institute, Department of Medical Sciences, Ministry of Public Health, Nonthaburi, Thailand; iMed.ULisboa-Institute for Medicines Research, Faculty of Pharmacy, University of Lisbon, Lisbon, 1649004, Portugal; Faculty of Epidemiology and Population Health, London School of Hygiene & Tropical Medicine, WC1E 7HT London, UK

**Keywords:** Amplicon sequencing, Amplicon design, *Mycobacterium tuberculosis*, tuberculosis, sequencing, genomics

## Abstract

Amplicon sequencing (Amp-Seq) of *Mycobacterium tuberculosis* genes associated with drug resistance and strain typing offers a cost-effective approach for profiling infections and tailoring the clinical management of tuberculosis. However, Amp-Seq assays require continual updates to incorporate new loci and mutations linked to drug resistance. We introduce TOAST (Tuberculosis Optimised Amplicon Sequencing Tool), a customisable software tool that optimises amplicon design across any loci and sequencing platforms (e.g., Illumina, Oxford Nanopore Technology (ONT)), informed by an integrated and expanding database of mutations from >50K *M. tuberculosis* isolates. TOAST software allows users to define parameters such as melting temperature, amplicon length, and GC content, while accounting for potential primer interactions like homodimer formation and non-specific binding. To demonstrate its robustness, we designed 33 amplicons in a single multiplex group, prioritising coverage of resistance-associated mutations in the form of insertions, deletions and single nucleotide polymorphisms (SNPs) for 13 different drugs. An efficient experimental protocol was established, resulting in a minimum depth coverage exceeding 50-fold for each amplicon region as validated by ONT sequencing of two clinical samples with multi-drug resistance. TOAST software enables the development of Amp-Seq assays for the rapid detection of drug-resistant TB, enhancing treatment strategies, improving outcomes, and curbing resistant infections. The cost-effectiveness and adaptability of Amp-Seq approaches make it crucial for managing TB in resource-limited settings, advancing global control efforts.

## Introduction

Tuberculosis (TB), caused by *Mycobacterium tuberculosis* (Mtb), remains a significant global public health challenge, with an estimated 10.8 million new infections and 1.3 million deaths reported in 2023^1^. The emergence of drug-resistant Mtb has further exacerbated this public health crisis. Worldwide, an estimated 3.2% of newly diagnosed TB cases and 16% of cases with prior treatments are resistant to rifampicin (RR-TB) or in addition to isoniazid (HR-TB), together referred to as multidrug-resistant TB (MDR-TB). Furthermore, a significant subset (6.2%) of these MDR-TB/RR-TB cases escalate to extensively drug-resistant TB (XDR-TB), showing additional resistance to fluoroquinolones and at least one Group A anti-TB drug, such as bedaquiline or linezolid^1^. Addressing the global health challenge of drug-resistant TB necessitates the development of rapid, accurate, and cost-effective diagnostic tools for Mtb detection and the subsequent identification of drug resistance.

The resistance to drugs is primarily caused by genetic mutations affecting either the drug’s target genes or the genes responsible for activating enzymes. Such mutations are usually SNPs, although small insertions and deletions (indels), as well as loss of function variants and large deletions capable of eradicating entire genes, are also known to induce resistance^2^. Around ∼50 loci in Mtb genome (total size 4.4 Mb; ∼4,000 genes) have been linked to drug resistance across 13 anti-TB drugs ^3^. Knowing the complete set of mutations driving drug resistance can inform diagnostic design^4^. Recent data from next-generation sequencing platforms have enabled the identification of circulating resistance mutations and the discovery of novel ones through phenotypic-genotypic association analyses. Approaches such as genome-wide association studies (GWAS), machine learning, and phylogenetic methods have played a pivotal role in these advancements ^5 6–9^. Complementary informatics tools, such as TB-Profiler^3^, have been developed to predict genotypic drug resistance from sequence data^5^, and are likely to replace laboratory-based phenotypic testing that can be time inefficient.

Whole genome sequencing (WGS) of Mtb has gained traction for both clinical and epidemiological investigations, including through the implementation of Illumina and Oxford Nanopore Technology (ONT) platforms. To improve cost-effectiveness for low resource settings, it is possible to target a high number of genes (*e*.*g*., drug resistance loci) across many samples using an amplicon-based approach (Amp-Seq) on NGS platforms or focus on the multiplexing of whole genomes if transmission is important. Unlike WGS, Amp-Seq selectively amplifies target sequences using Polymerase Chain Reaction (PCR) before high-throughput sequencing. The design of amplicon primers is a pivotal step in determining the accuracy and efficiency of the subsequent sequencing, including the standardisation of melting temperatures (Tm) to avoid non-specific binding or inefficient amplification, compromising the integrity of Amp-Seq. Further, structural considerations in primer design, such as avoiding homopolymers, hairpins, homodimers, and heterodimers, are crucial to the optimisation of Amp-Seq assays.

Here we introduce the TOAST (Tuberculosis Optimized Amplicon Sequencing Tool) software tool aimed at addressing the challenges in amplicon primer design for TB sequencing leveraging the package Primer3 software^10^. TOAST optimises for Tm, homopolymers, hairpins, homodimers considerations, and through the application of an in-house pipeline, avoids heterodimer and alternative binding. Our automated tool uses a growing library of Mtb mutations to ensure users can design primers that target and capture important variants. The robustness of primer sets for TB is demonstrated on drug-resistant Mtb from Portugal, and the software is publicly available (https://github.com/linfeng-wang/TOAST). A website-based version is also available (https://genomics.lshtm.ac.uk/TOAST).

## Results

### TOAST software

The tool, implemented in Python, is connected to a database comprising 50,723 Mtb isolates with whole-genome sequencing (WGS) data (the ‘50K dataset’) ^6,8^. This dataset represents the major Mtb lineages: L1 (9.1%), L2 (27.6%), L3 (11.8%), and L4 (48.3%). Among these, 9,546 (29.2%) exhibited genotypic HR-TB, 7,974 (24.4%) RR-TB, and 5,385 (16.5%) showed at least MDR-TB.

### Amplicons and mutations covered

Using TOAST, thirty-three amplicon primers (forward and reverse pairs) were designed in 50 bp increments, ranging from 300 bp to 800 bp, with each increment repeated three times. These primers covered 2,377 known mutations across 25 genes associated with drug resistance, selected based on mutation frequency observed in the 50K dataset (**Figure 1, Table S1**). The choice of 33 amplicons maximised mutation coverage, minimised diminishing returns on covering isolated rare mutations, and avoided primer clashes associated with higher numbers of amplicons (**Figure S1**). This set of ONT-sized amplicons (500bp), designed by TOAST, achieved 99.1% coverage of the established mutation loci. When tested with Illumina-sized amplicons (300bp), the coverage reached 97.0%. On the other hand, longer amplicons (800 bp) suitable for ONT can also be generated, potentially expanding the flexibility of experimental designs and enabling the capture of more complex variants.

**Figure 1.**
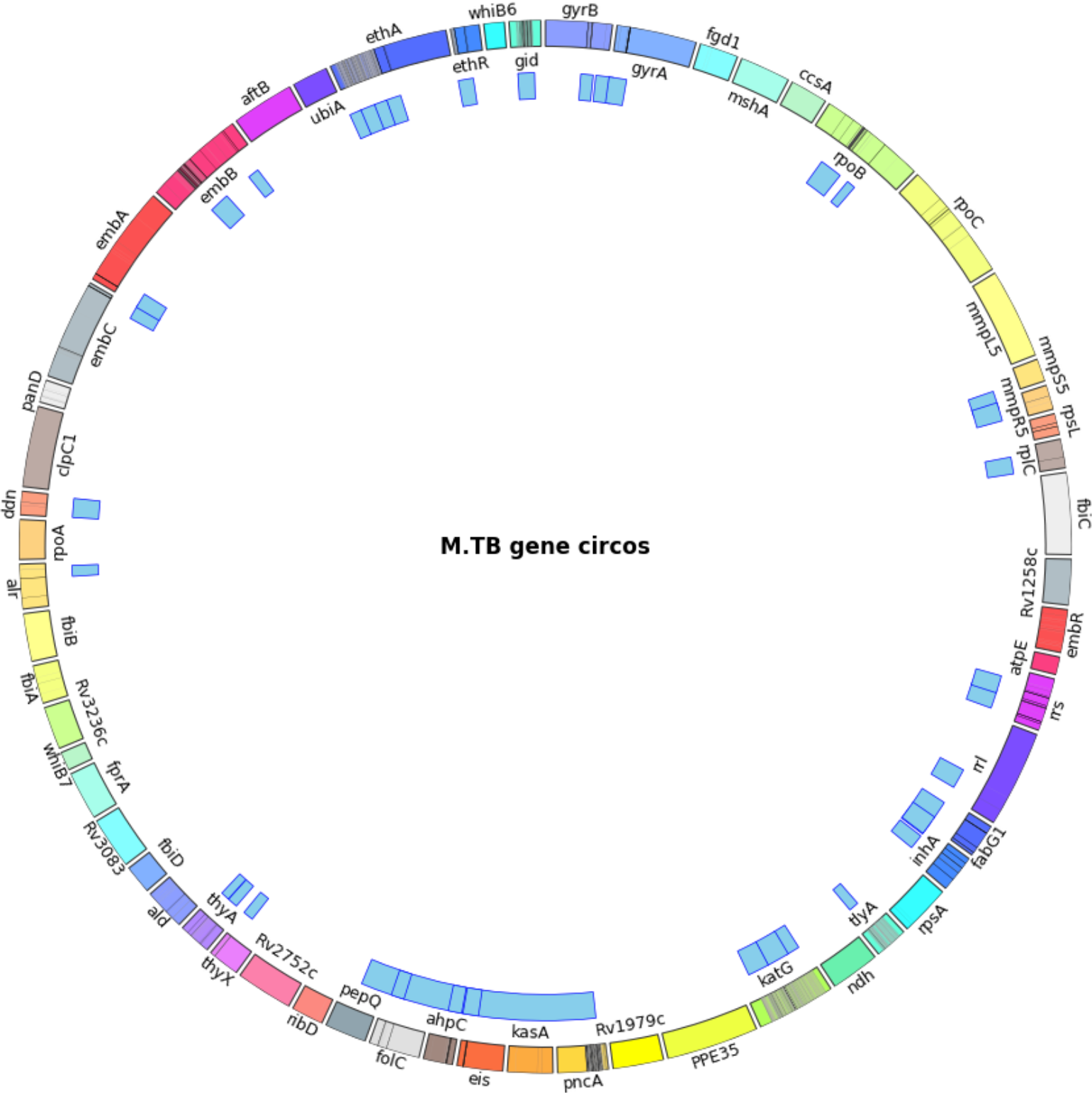
The 33 amplicons across the *M. tuberculosis* genome. Mtb gene feature (multi-coloured outer ring) with mutation cytobands (more mutation with higher frequencies is in deeper colour) and amplicon position (Blue markers).

**Figure 2.**
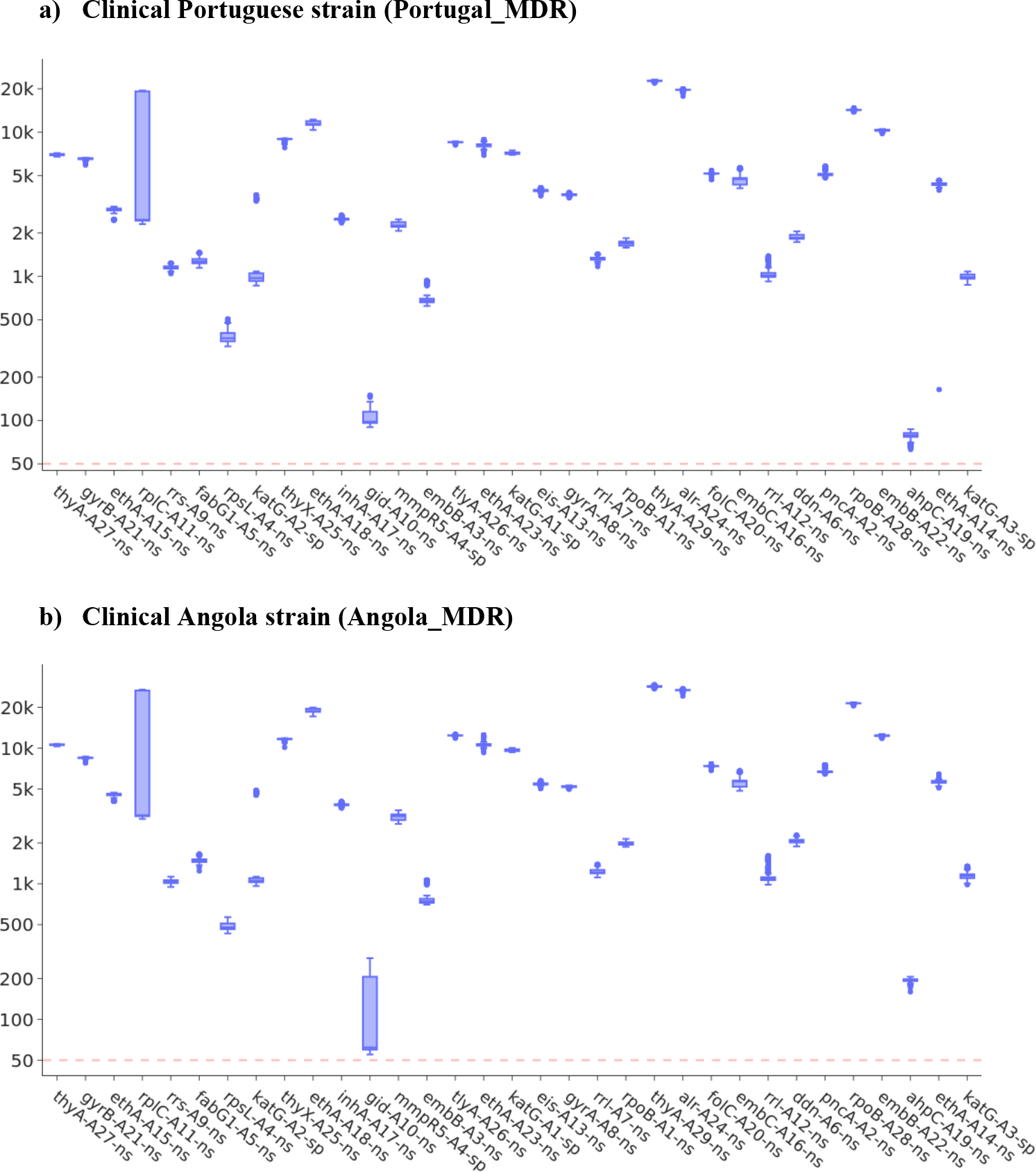
Boxplots of ONT sequence coverage depth for the 33 amplicons. Dashed red line marks a 50-fold coverage threshold

The amplicon coverage specifically included *katG* (isoniazid, 6 amplicons) and *mmpR5* (bedaquiline, 1 amplicon) in which loss of function resistance associated mutations can occur at any position. The remaining genes covered resistance to ethionamide (*ethA*, 4 amplicons), PAS (*folC, thyX, thyA*; 4), ethambutol (*embB*, 3), aminoglycosides/tuberactinomycins (*rrs*, 2), streptomycin (*rpsL, gid*; 2), linezolid (*rrl, rplC*; 2), rifampicin (*rpoB*, 2), fluoroquinolones (gyrA, *gyrB*, 2), capreomycin (*tlyA*, 1), D-cycloserine (*alr*, 1), kanamycin (*eis*, 1), pretomanid (*ddn*, 1) and pyrazinamide (*pncA*, 1). All drug resistance genes were completely covered by the designed amplicons, except for several loci with known localised genetic regions containing all or most of the known resistance-linked mutations: *gid* (99%), *rpoB* (98%), *katG* (92%), *embB* (92%), *ahpC* (92%), *inhA* (88%), *pncA* (72%), *embA* (50%), and *thyA* (29%). Importantly, the amplicons successfully captured most of the WHO-defined resistance mutations^11^ (742/764, 97.1%), with exceptions observed in *pncA* (155/166, 93.4%) for pyrazinamide resistance and *tlyA* (19/30, 63.3%) for capreomycin resistance. The *pncA* gene harbours a high number of low-frequency mutations and tends to receive lower coverage due to the weighting scheme based on sample frequency from the 50K database. However, the weighting scheme for mutation coverage can be adjusted to prioritise enhanced coverage in user-specified regions, providing flexibility for targeted applications while enabling the development and implementation of locally adapted testing. This comprehensive amplicon design ensures robust coverage of known resistance loci, supporting detailed genomic surveillance and the detection of emerging drug-resistant mutations.

### Validation

The amplicons were validated through ONT sequencing of Mtb isolate DNA sourced from clinical MDR cases in Portugal (Portugal_MDR, 272439 reads; median length 401 bps) and Angola (Angola_MDR; 394977 reads; median length 406 bps). The vast majority of ONT reads mapped to the targeted 25 loci for both Portugal_MDR (261503/272439; 96.0%) and Angola_MDR (374609/394977; 94.8%). Most amplicon regions had high depth (>500-fold), except for *gid* (A10-ns) and *ahpC* (A19-ns) amplicons (**S1 Table**). The minimum depth of coverage for *gid* A10-ns was 63-fold (Portugal_MDR 90-fold; Angola_MDR 63-fold) and for *ahpC* A19-ns was 55-fold (Portugal_MDR 55-fold; Angola_MDR 160-fold). The drug resistance profiles obtained from amplicon sequencing corresponds perfectly to that obtained from WGS^12^.

The 33 amplicons ranged in size from 300 to 800 bp, with sequencing depth decreasing as amplicon size increased, likely due to reduced amplification efficiency (**S2 Figure**). Nevertheless, even amplicons exceeding 900 bp achieved a sequencing depth greater than 50-fold in our validation run (**S2 Figure**). While an inverse relationship between amplicon size and depth was observed, individual primer designs significantly influenced the resulting depth, occasionally causing substantial drops. This effect was more pronounced for shorter amplicons, whereas longer amplicons exhibited more consistent depth values.

## DISCUSSION

The TOAST tool integrates the Primer3 primer design software with a large and growing collection of WGS data, currently consisting of >50K clinical Mtb isolates^6,8^. It enables automated amplicon design optimised to cover the most frequently occurring resistance mutations. TOAST is highly customisable, allowing users to input their own list of mutations and loci, and to expand the amplicon set to include emerging drug resistance targets without requiring extensive reconfiguration. To demonstrate its utility, we used TOAST to design a 33-amplicon set targeting drug resistance genes, similar to previously proposed and commercially available amplicon sets^13,14 15^. However, TOAST is not limited to drug resistance targets; it can also be tailored to identify signals of particular strain types or transmitting clones^16^, as well as the presence of *pe/ppe* genes, which may serve as vaccine targets^17^.

Integrating TOAST with emerging sequencing technologies holds significant potential for public health, disease tracking, and diagnostics. Its ability to rapidly design and deploy customised amplicon sets enhances the detection of drug-resistant strains, enabling timely treatment decisions and containment strategies. Beyond TB, TOAST can be adapted to other bacterial species by incorporating relevant genomic data and target regions. This flexibility makes TOAST a valuable tool in microbial genomics, where rapid adaptation to emerging threats is essential. By keeping pace with the dynamic genetic landscape of bacterial pathogens, TOAST contributes to improved surveillance, diagnostics, and disease management, ultimately supporting better patient outcomes and global health initiatives.

## Methods

### Software overview

TOAST (Tuberculosis Optimized Amplicon Sequencing Tool) is a command-line program designed to generate and identify TB amplicons based on user-defined parameters, such as the number and size of amplicons, while accommodating predefined existing amplicons. An iterative mutation search algorithm systematically scans a user-provided database^3^ within a defined window size (amplicon size), positioning amplicons at locations with the highest priority scores. Once optimal positions are identified, Primer3 methodology is employed to design primers within an extended range of the selected windows, ensuring they meet predefined criteria for melting temperature, GC content, hairpin prevention, and homodimer formation. The resulting primers are then screened in an all-versus-all manner to check for heterodimer binding and alternative binding at unintended genomic locations (**S3 Figure**).

Using these principles, TOAST can also estimate the number of amplicons required to achieve near-complete genome coverage (e.g., 99%) for a given amplicon size. The tool provides a detailed output, including primer sequences, positions, and genomic regions covered. Outputs can be visualized using tools such as the Interactive Genomic Viewer (IGV) ^18^ (**S4 Figure**) alongside mapped sequencing data files (e.g., bam format), enabling seamless integration and analysis. This automated and robust design pipeline streamlines TB amplicon sequencing, making it a versatile tool for targeted mutation analysis and genomic surveillance.

### Amplicon design

An optional preliminary step in TOAST involves estimating the number of amplicons required to cover all mutations in the provided database using the TOAST amplicon_no script. This step generates a line plot showing diminishing returns of mutation coverage as the number of amplicons increases. Following this, the TOAST design function can be used to generate detailed amplicon designs. Users can define the amplicon size (−a) and specify genes that must be fully covered (−sg). Nearly all design process variables are user-adjustable, while non-gene-specific runs require users to input the desired number of non-gene-specific amplicons (−nn). To facilitate primer amplification quality checks via gel electrophoresis, TOAST offers the -seg function, which allows the design of varying-sized amplicons. This function accepts comma-separated inputs for minimum amplicon size, maximum amplicon size, step size, and the number of amplicons per step. The algorithm then identifies the optimal combination of amplicons to cover each specific gene with minimal overlap. Remaining amplicons are assigned to non-specific regions based on mutation weighting, ordered from the largest to the smallest amplicon size.

By default, TOAST assumes the use of Q5 High-Fidelity DNA Polymerase. However, design criteria such as ideal melting temperature (Tm), GC content, primer size, mononucleotide repeats, self/pair complementarity, and DNA concentration can be customised through a user-defined JSON file using the -set option. Primer design leverages Primer3 software, which calculates a penalty score for each candidate primer. Primer3 methods determine the optimal primer pair for PCR by evaluating factors such as melting temperature, GC content, primer length, and self-complementarity.

Primers that fail to meet strict thresholds—such as maximum GC content or minimum melting temperature—are excluded. For remaining primers, penalty scores are computed based on weighted deviations from optimal values, factoring in end stability and potential mispriming. Primer pairs are additionally penalized for complementarity and melting temperature mismatches. The pipeline further evaluates primer pairs for alternative genomic binding and homodimer formation, discarding pairs that fail these checks, prioritizing those with the lowest penalty scores.

To ensure robust primer placement, a modifiable extended region—by default set to one-sixth of the amplicon size—is added to both ends of the amplicon. This buffer region maximises the chances of Primer3 locating suitable primer binding sites while preserving the amplicon ranges determined by the iterative mutation search algorithm. Primer sequences may include degenerate base pairs to account for mutations within binding sites, as identified from the 50K dataset.

The output includes a CSV file (Primer_design) containing primer details such as amplicon ID, primer ID, melting temperature, sequences, and genomic coordinates. A mutation inclusion report (mutation_inclusion) specifies which mutations are covered by each amplicon. TOAST also provides plotting functions to visualise amplicon distribution by gene, along with a comparison to reference amplicon designs if available. Additionally, a .bed file is generated, detailing amplicon positions, actual amplicon coverage, and primer coordinates, which can be visualized using the IGV.

### Culture, DNA extraction and sequencing

All primers were combined into a single pool for the amplification. A PCR master mix was prepared to a final volume of 25 μl, comprising 5 μl of Q5 Reaction Buffer (1X), 0.5 μl of dNTPs, 5 μl of High GC Enhancer, 0.25 μl of Q5 enzyme. 0.1μl of each primer, 1.5 μl of DNA sample. The rest is made up with miliQ water. The PCR cycling conditions were as follows: Tubes were placed in a thermal cycler set to an initial denaturation step of 30 seconds at 98°C, followed by 35 cycles of denaturation at 98°C for 10 seconds, annealing at 58°C for 40 seconds, and extension at 72°C for 55 seconds. After the cycles, a further extension at 72°C for 5 minutes was performed. After the PCR reaction, the products were checked with gel electrophoresis, confirming that the intended segments had been successfully generated (**S5 Figure**). Next, using this full set of primers, we conducted simultaneous amplifications on two MDR Mtb sourced from TB clinical cases in Portugal (Portuguese_MDR) and Angola (Angola_MDR) ^12^.

### Library Preparation and Amplicon Sequencing

The amplicon products were quantified using a DS-11 FX+ Spectrophotometer/Fluorometer (DeNovix) along with the Qubit™ 1X dsDNA High Sensitivity Assay Kit. Following the manufacturer’s protocol for ligation sequencing of amplicons, the amplicon library was prepared using the Native Barcoding Kit 96 V14 (SQK-NBD114.96, ONT). The prepared library was subsequently sequenced on the MinION Mk1B platform using R10.4.1 flow cells (ONT) and operated with MinKNOW v24.06.14 software (ONT). The Dorado baseballer was used to process the resulting data.

## Supporting information

Designed primers

## Data availability

Raw sequencing data generated in this project is available from the ENA archive (accession numbers: PRJEB84199). The primers designed and tested in this study can be found in the supplementary data.

## Code availability

The code can be found on github (https://github.com/linfeng-wang/TOAST). It is also available as a package (https://genomics.lshtm.ac.uk/TOAST).

## Acknowledgements

LW is funded by a BBSRC LIDO studentship (Ref. BB/T008709/1). NT is funded by a Thailand government, Ministry of Public Health PhD scholarship. TGC and SC are funded by the UKRI (BBSRC BB/X018156/1; MRC MR/R020973/1, MRC MR/X005895/1; EPSRC EP/Y018842/1)). NT, WS and SM were funded by the Department of Medical Sciences (Thailand Ministry of Public Health) and the Thailand Health Systems Research Institute (HSRI 64-179). The funders had no role in study design, data collection and analysis, decision to publish, or preparation of the manuscript. The authors declare no conflicts of interest.

## Author contributions

SC, TGC and JEP conceived and directed the project. LW, NT, JP and SC undertook sample processing and DNA extraction. JP performed bacterial culture. MT and SC performed sequencing. LW developed the software under the supervision of SC, TGC and JEP. JT and MH contributed bioinformatic tools. WS, SM and NT contributed reagents. LW, SC, TGC and JEP interpreted the results. LW wrote the first draft of the manuscript. All authors commented and edited various versions of the draft manuscript and approved the final manuscript. LW, TGC and JEP compiled the final manuscript.

