## Supplementary for "A novel tool for designing targeted gene amplicons and an optimised set of primers for high-throughput sequencing in tuberculosis genomic studies"

**S1 Table**
**Amplicons and genes covered**

| Amplicon ID | Gene | Drug | No. mutations* | Median Portugal_MDR coverage** | Median Angola_MDR coverage*** |
|---|---|---|---|---|---|
| A1-ns | *rpoB* | Rifampicin | 160 | 1693 | 1969 |
| A1-sp | *katG* | Isoniazid | 116 | 7148 | 9642 |
| A2-sp | *katG* | Isoniazid | 296 | 972 | 1052 |
| A2-ns | *pncA* | Pyrazinamide | 961 | 5119 | 6681 |
| A3-sp | *katG* | Isoniazid | 308 | 987 | 1145 |
| A3-ns | *embB* | Ethambutol | 89 | 678 | 745 |
| A4-sp | *mmpR5* | Bedaquiline | 6 | 2257 | 3147 |
| A4-ns | *rpsL* | Streptomycin | 11 | 373 | 477 |
| A5-ns | *fabG1* | Isoniazid | 9 | 1269 | 1479 |
| A6-ns | *ddn* | Pretomanid | 3 | 1852 | 2045 |
| A7-ns | *rrl* | Linezolid | 2 | 1328 | 1204 |
| A8-ns | *gyrB* | Fluoroquinolones | 17 | 3687 | 5197 |
| A9-ns | *rrs* | Aminoglycosides | 6 | 1144 | 1038 |
| A10-ns | *gid* | Streptomycin | 136 | 98 | 62 |
| A11-ns | *rplC* | Linezolid | 1 | 2461 | 3190 |
| A12-ns | *rrs* | Aminoglycosides | 11 | 1016 | 1083 |
| A13-ns | *eis* | Kanamycin | 10 | 3934 | 5428 |
| A14-ns | *ethA* | Ethionamide | 357 | 4359 | 5642 |
| A15-ns | *ethA* | Ethionamide | 98 | 2885 | 4511 |
| A16-ns | *embA* | Ethambutol | 6 | 4722 | 5688 |
| A17-ns | *inhA* | Isoniazid | 7 | 2488 | 3814 |
| A18-ns | *ethA* | Ethionamide | 242 | 11324 | 18630 |
| A19-ns | *ahpC* | Isoniazid | 14 | 79 | 195 |
| A20-ns | *folC* | PAS | 16 | 5179 | 7366 |
| A21-ns | *gyrB* | Fluoroquinolone | 24 | 6582 | 8503 |
| A22-ns | *embB* | Ethambutol | 4 | 10291 | 12330 |
| A23-ns | *ethA* | Ethionamide | 83 | 8156 | 10618 |
| A24-ns | *alr* | D-cycloserine | 2 | 19676 | 26800 |
| A25-ns | *thyX* | PAS | 1 | 8994 | 11694 |
| A26-ns | *tlyA* | Capreomycin | 149 | 8532 | 12413 |
| A27-ns | *thyA* | PAS | 8 | 7020 | 10617 |
| A28-ns | *rpoB* | Rifampicin | 1 | 14252 | 21415 |
| A29-ns | *thyA* | PAS | 11 | 22726 | 28514 |

PAS Para-aminosalicylic acid; * in the 50K dataset; ** ONT sequencing of XXXX; *** ONT sequencing of the Clinical Angola strain

**S2 Table**

**Coverage of the 33 amplicons of the WHO catalogue of drug resistance mutation**

| Drug | No. detected mutations | WHO mutations | % covered |
|---|---|---|---|
| Ethionamide | 270 | 270 | 100 |
| Pyrazinamide | 155 | 166 | 93 |
| Streptomycin | 143 | 143 | 100 |
| Isoniazid | 97 | 97 | 100 |
| Rifampicin | 45 | 45 | 100 |
| Capreomycin | 19 | 30 | 63 |
| Kanamycin | 8 | 8 | 100 |
| Aminoglycosides | 4 | 4 | 100 |
| Ethambutol | 1 | 1 | 100 |

**S1 Figure**

**Coverage of mutations identified in the candidate genes in the 50K dataset across the number of amplicons**

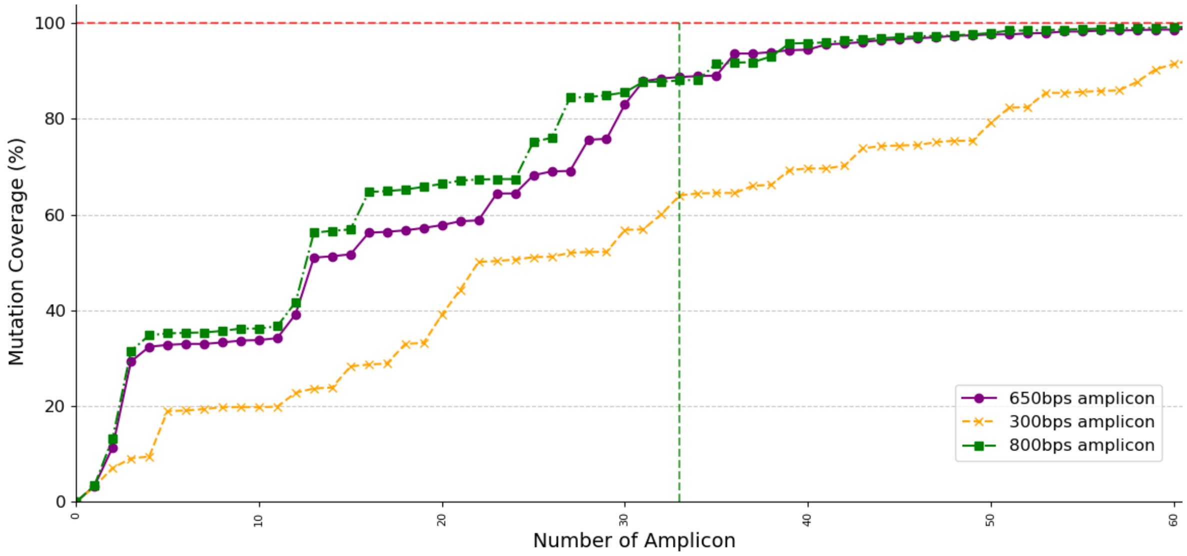


Red dashed line: 100% mutation coverage threshold. Green dashed line: 33 amplicons

### S2 Figure

### The distribution of coverage depth across amplicon sizes

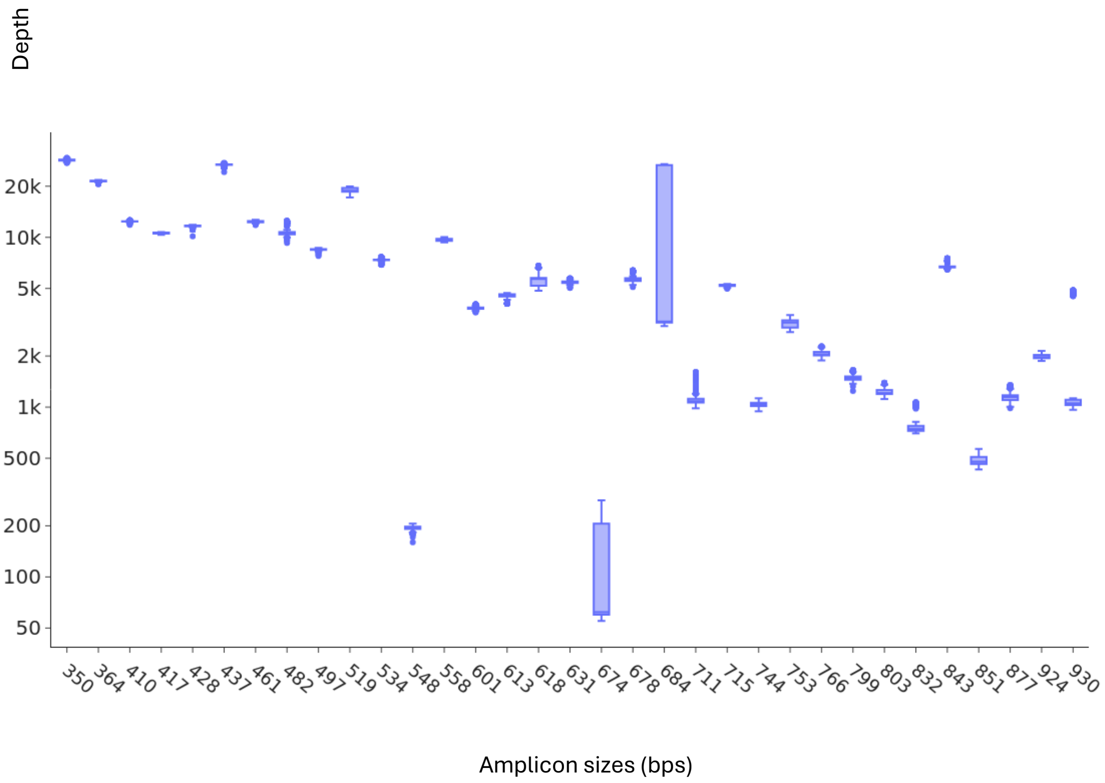

**S3 Figure**

**An overview of the TOAST workflow**

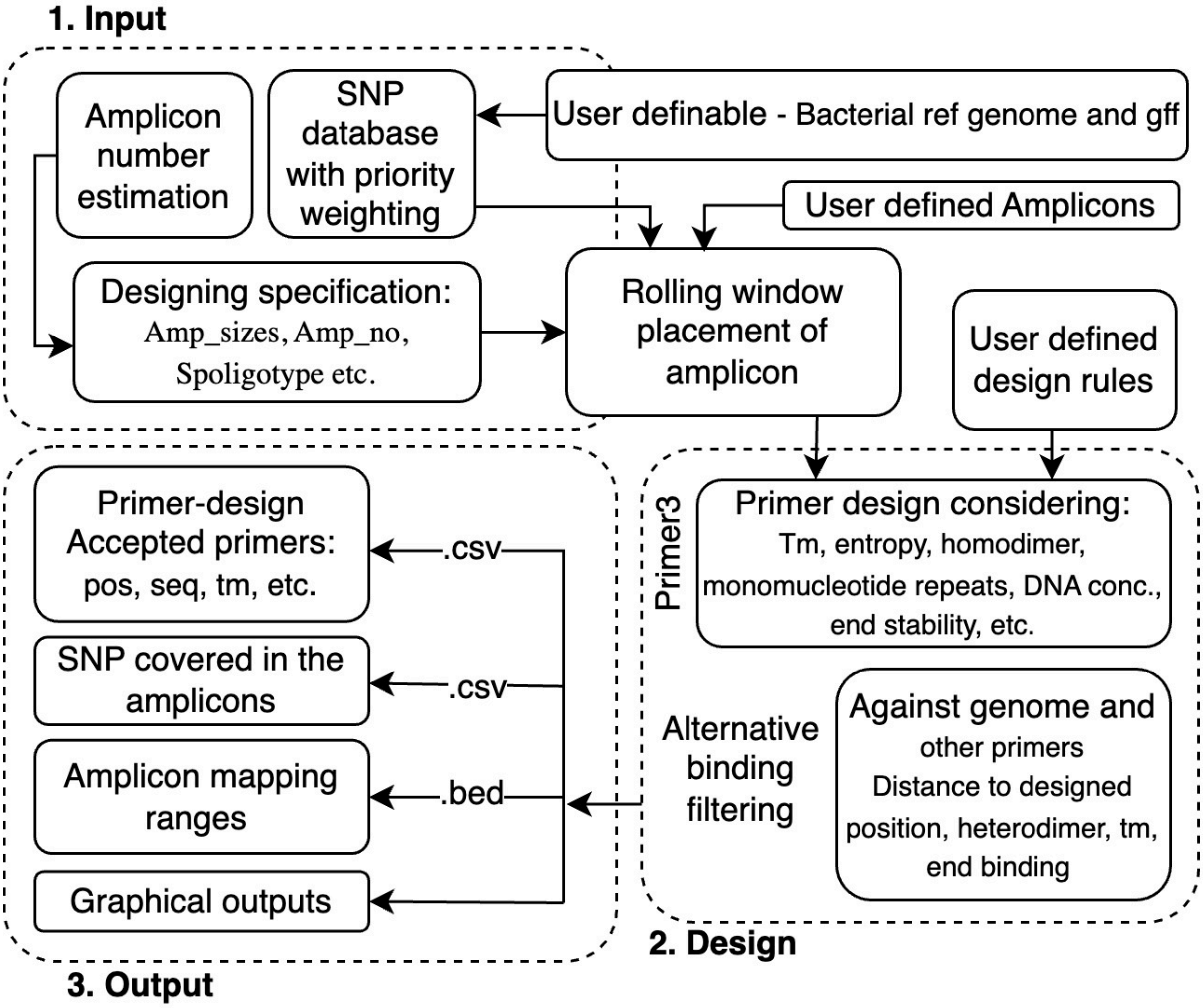


The TOAST workflow comprises three stages. In the input stage, mutation priority weighting (from a 50k dataset) and user-defined parameters guide amplicon placement using a rolling window approach. In the design stage, primers are generated using Primer3, considering factors such as Tm, entropy, and stability, followed by in-house filtering to eliminate alternative binding sites. The output stage provides primers with low penalty values and no alternative binding, along with SNP coverage details, amplicon mapping ranges, and graphical outputs.

**S4 Figure**

**IGV visualisation of designed amplicons**

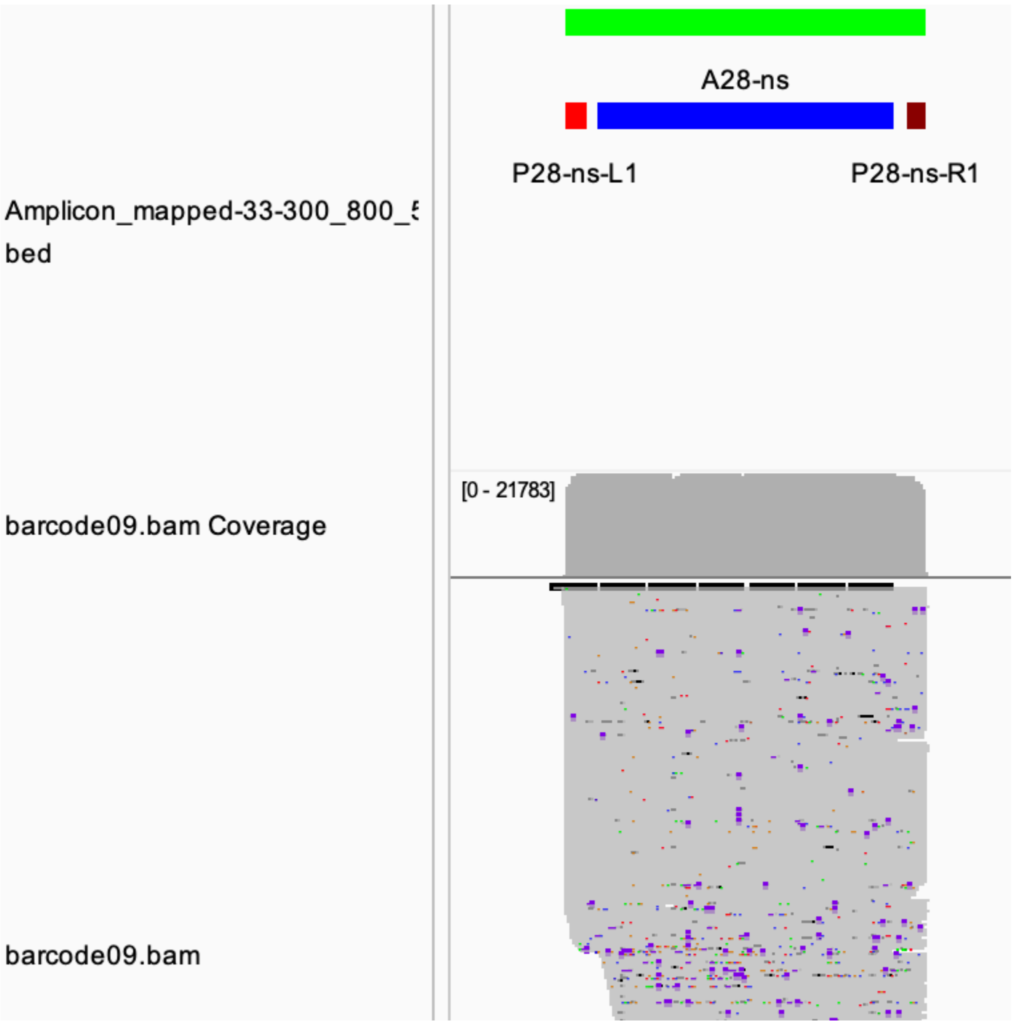


Red: left primer; brown: right primer; green: total amplicon size with buffer region and primer; blue: aimed range for amplification without buffer region and primer; grey: read coverage.

**S5 Figure**

**Gel electrophoresis photo for the 33 amplicon PCR.**

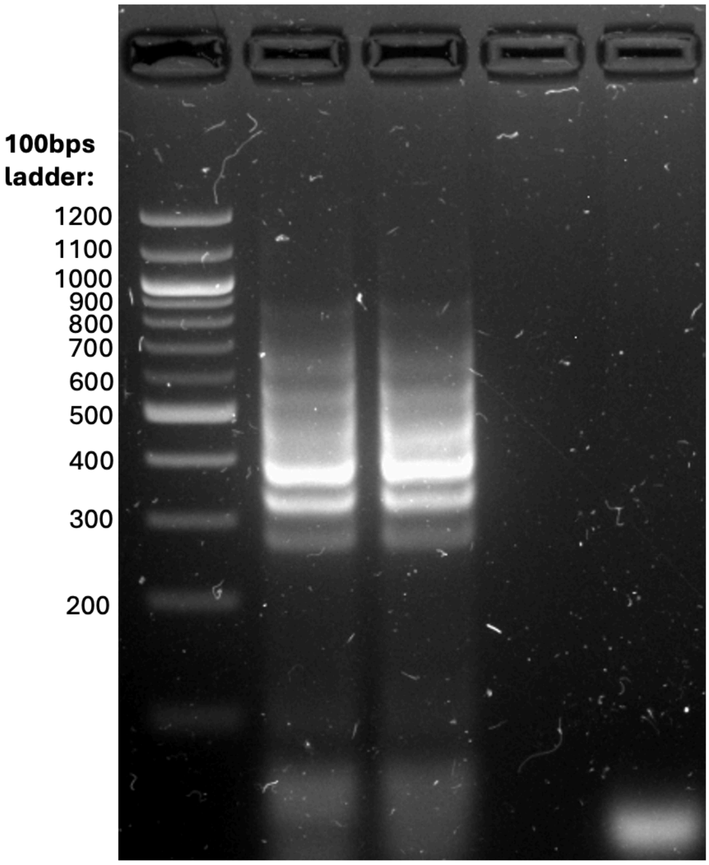


First column 58C, Second column 60C. Third column: gap, fourth column: water
